# Functional plasticity coupled with structural predispositions in auditory cortex shape successful music category learning

**DOI:** 10.1101/2021.05.12.443818

**Authors:** Kelsey Mankel, Utsav Shrestha, Aaryani Tipirneni-Sajja, Gavin M. Bidelman

## Abstract

Categorizing sounds into meaningful groups helps listeners more efficiently process the auditory scene and is a foundational skill for speech perception and language development. Yet, how auditory categories develop in the brain through learning, particularly for nonspeech sounds, is not well understood. Here, we asked musically naïve listeners to complete a brief (∼20 min) training session where they learned to identify sounds from a nonspeech continuum (minor-major 3rd musical intervals). We used multichannel EEG to track behaviorally relevant neuroplastic changes in the auditory event-related potentials (ERPs) pre- to post-training. To rule out mere exposure-induced changes, neural effects were evaluated against a control group of 14 nonmusicians who did not undergo training. We also compared individual categorization performance with structural volumetrics of bilateral primary auditory cortex (PAC) from MRI to evaluate neuroanatomical substrates of learning. Behavioral performance revealed steeper (i.e., more categorical) identification functions in the posttest that correlated with better training accuracy. At the neural level, improvement in learners’ behavioral identification was characterized by smaller P2 amplitudes at posttest, particularly over right hemisphere. Critically, learning-related changes in the ERPs were not observed in control listeners, ruling out mere exposure effects. Learners also showed smaller and thinner PAC bilaterally, indicating superior categorization was associated with structural differences in primary auditory brain regions. Collectively, our data suggest successful auditory categorical learning of nonspeech sounds is characterized by short-term functional changes (i.e., greater post-training efficiency) in sensory coding processes superimposed on preexisting structural differences in bilateral auditory cortex.

## INTRODUCTION

Classifying continuously varying sounds into meaningful categories like phonemes or musical intervals enables more efficient processing of an auditory scene (Bidelman et al., 2020). Categorization of auditory stimuli is also a foundational skill for language development and is believed to arise from both learned and innate factors (Livingston et al., 1998; Mankel, Barber, et al., 2020; Mankel, Pavlik Jr, et al., 2020; Perez-Gay Juarez et al., 2019; Rosen & Howell, 1987). Auditory categories are further shaped by experiences such as speaking a second language (Escudero et al., 2011; Lively et al., 1993; Perrachione et al., 2011) or musical training (Bidelman & Walker, 2019; Bidelman et al., 2014; Wu et al., 2015), suggesting flexibility in categorical perception with learning. While the behavioral aspects of category acquisition are well documented, the underlying neural mechanisms and the influence of individual differences in shaping this process are poorly understood.

Characterizing the neurobiology of category acquisition is typically confounded by prior language experience and the overlearned nature of speech (Liu & Holt, 2011). For example, perceptual interference from native-language categories can impede the learning of foreign speech sounds (Flege & MacKay, 2004; Francis et al., 2008; Guion et al., 2000). Instead, nonspeech stimuli (e.g., music) offers the ability to probe the neural mechanisms of nascent category learning without the potential confounds of language background or automaticity that stems from using speech materials (Goudbeek et al., 2009; Guenther et al., 1999; Liu & Holt, 2011; Smits et al., 2006; Yi & Chandrasekaran, 2016). In this regard, musical categories (i.e., intervals, chords) offer a fresh window into tabula rasa category acquisition. Indeed, nonmusicians are unable to adequately categorize musical stimuli despite their exposure to music in daily life (Bidelman & Walker, 2019; Howard et al., 1992; Klein & Zatorre, 2011; Locke & Kellar, 1973; Siegel & Siegel, 1977). While several studies have assessed category learning of musical intervals, they either used highly trained listeners (Burns & Ward, 1978) or focused on different training methods that maximize learning gains (Little et al., 2019; Pavlik Jr et al., 2013). To our knowledge, no study has assessed the *neural* changes associated with category learning in music.

Speech categorization is believed to emerge in the brain by around N1 of the cortical event-related potentials (ERPs) and fully manifests by P2 (i.e., ∼150-200 ms; Alho et al., 2016; Bidelman & Lee, 2015; Bidelman, Moreno, et al., 2013; Bidelman & Walker, 2017; Mankel, Barber, et al., 2020; Ross et al., 2013). Fewer studies have examined the electrophysiological underpinnings of music categorization, but evidence from musicians suggests a similar neural time course (Bidelman & Walker, 2019). Functional magnetic resonance imaging suggests that categorization training leads to a decrease in perceptual sensitivity for within-category stimuli in auditory cortex while learning to discriminate categorical sounds shows the opposite effect— greater sensitivity to differences between stimuli (Guenther et al., 2004). Still, the majority of studies on category learning have involved speech. Although there are probably some parallels (Liu & Holt, 2011), it remains unclear whether the neuroplastic changes that arise when rapidly learning nonspeech categories (e.g., music) parallels that of speech.

More generally, auditory perceptual learning studies have reported changes in both early sensory-evoked (i.e., N1, P2) and late slow-wave ERP responses following training (Alain et al., 2010; Alain et al., 2007; Atienza et al., 2002; Ben-David et al., 2011; Bosnyak et al., 2004; Carcagno & Plack, 2011; Tong et al., 2009; Tremblay et al., 2001; Tremblay & Kraus, 2002; Tremblay et al., 2009; Wisniewski et al., 2020). A true biomarker of learning, however, should vary with learning performance (Tremblay et al., 2014). Because modulations in P2 amplitudes can occur with mere passive stimulus exposure in the absence of training improvements, some posit P2 reflects aspects of the task acquisition process rather than training or perceptual learning, *per se* (Ross et al., 2013; Ross & Tremblay, 2009; Tremblay et al., 2014). Given the equivocal role of P2 in relation to auditory learning, we aimed to re-adjudicate whether changes in P2 scale with individual behavioral outcomes as listeners rapidly acquire novel music categories.

There is also significant variability in the acquisition of auditory categories (e.g., Golestani & Zatorre, 2009; Howard et al., 1992; Mankel, Pavlik Jr, et al., 2020; Silva et al., 2020), especially for speech (Díaz et al., 2008; Fuhrmeister & Myers, 2021; Kajiura et al., 2021; Mankel, Barber, et al., 2020; Wong et al., 2007). More successful learners show greater neural activation, particularly in auditory cortex (Díaz et al., 2008; Kajiura et al., 2021; Wong et al., 2007). Such variability might be attributable to differences in the creation or retrieval of long-term memories for prototypical vs. non-prototypical sounds during learning (Golestani & Zatorre, 2009). However, we have previously shown better categorizers show efficiencies even in early sensory processing (∼150-200 ms), suggesting stimulus representations themselves are tuned at the individual level rather than later memory-related processes, *per se* (Mankel, Barber, et al., 2020).

In addition to differences in functional processing, individual categorization abilities may be partially driven by preexisting structural advantages within the brain (Fuhrmeister & Myers, 2021). Paralleling the left hemisphere bias for speech (Binder et al., 2004; Bouton et al., 2018; Lee et al., 2012; Myers et al., 2009), categorization of musical sounds is believed to involve a frontotemporal network in the right hemisphere, including key brain regions such as the primary auditory cortex (PAC), superior temporal gyrus (STG), and inferior frontal gyrus (IFG) (Bidelman & Walker, 2019; Klein & Zatorre, 2011, 2015; Mankel, Barber, et al., 2020). PAC/STG size (primarily right hemisphere) has also been associated with perception of relative pitch and musical transformation judgments (Foster & Zatorre, 2010), melodic interval perception (Li et al., 2014), spectral processing (Schneider et al., 2005), and even musical aptitude (Schneider et al., 2002). To our knowledge, few studies have examined the structural correlates of categorization differences on the individual level. In the domain of speech, faster, more successful learners of nonnative phonemes exhibit larger left Heschl’s gyrus (Golestani et al., 2007; Wong et al., 2008) and parietal lobe volumes (Golestani et al., 2002). Additionally, better and more consistent speech categorizers show increased right middle frontal gyrus surface area and reduced gyrification in bilateral temporal cortex (Fuhrmeister & Myers, 2021). We thus hypothesized that successful category learning in *music* would be predicted by neuroanatomical differences (e.g., gray matter volume, cortical thickness), with perhaps effects favoring right PAC.

The aim of this study was to examine the functional and structural neural correlates of auditory category learning following short-term identification training of music sound categories. Musical intervals allowed us to track sound-to-label learning without the potential lexical-semantic confounds inherent to using speech materials (Liu & Holt, 2011). We measured learning-related changes in the cortical ERPs in musically naïve listeners against a no-contact control group to determine the specificity of neuroplastic effects. If rapid auditory category learning is related to enhanced sensory encoding of sound, we predicted changes in early brain activity manifesting at or before auditory object formation (i.e., prior to ∼250 ms; P2). If instead, short-term learning is associated with later cognitive processes related to decision and/or task strategy, we expected neural effects to emerge later in the ERP time course (e.g., late slow waves > 400-500 ms; Alain et al., 2007). Additionally, we anticipated successful learners would recruit neural resources in right auditory cortices, mirroring the left hemispheric specialization supporting speech categorization (Bidelman & Walker, 2019; Joanisse et al., 2007; Klein & Zatorre, 2011; Liebenthal et al., 2005). Our findings show that successful auditory category learning is characterized by both structural and functional differences in right auditory cortex. The presence of anatomical differences along with ERP changes specific to learning suggest that the acquisition of auditory categories depend on a layering of preexisting and short-term plastic changes in brain function.

## MATERIALS & METHODS

### Participants

Our sample included N=33 participants. Nineteen young adults (16 females) participated in the training task. An additional fourteen (7 females) served as a control group (data from Mankel, Barber, et al., 2020). All had normal hearing (thresholds ≤25 dB SPL, 250-8000 Hz), were right-handed (Oldfield, 1971), and had no history of neurological disorders. Participants completed questionnaires that assessed education level, socioeconomic status (SES) (Entwislea & Astone, 1994), language history (Li et al., 2006), and music experience. Groups were comparable in age (learners: *µ* = 24.9 ± 4.0 yrs, controls: *µ* = 24.9 ± 1.7 yrs; *p* = 0.5492), education (learners: *µ* = 18.5 ± 3.3 yrs, controls: *µ* = 17.3 ± 3.0 yrs; *p* = 0.3211), and SES (rating scale of average parental education from 1 [some high school education] to 6 [PhD or equivalent]; learners: *µ* = 4.6 ± 1.3, controls: *µ* = 4.1 ± 0.6; *p* = 0.1085). All were fluent in English though six reported a native language other than English. We excluded tone language speakers as these languages improve musical pitch perception (Bidelman, Hutka, et al., 2013). To ensure participants were naïve to the music-theoretic labels for pitch intervals, we required participants have no more than three years total of formal music training on any combination of instruments and none within the past five years. Critically, groups did not differ in prior music training (learners: *µ* = 1.1 ± 1.0 yrs, controls: *µ*=0.6 ± 0.8 yrs; *p*=0.1446). All participants gave written informed consent according to protocol approved by the University of Memphis Institutional Review Board and were compensated monetarily for their time.

### Stimuli

We used a five-step musical interval continuum to assess category learning of non-speech sounds continuum (Bidelman & Walker, 2017; Mankel, Pavlik Jr, et al., 2020). Individual notes of each dyad were constructed of complex tones consisting of 10 equal amplitude harmonics added in cosine phase. Each token was 100 ms in duration with a 10 ms rise/fall time to reduce spectral splatter. The bass note was fixed at a fundamental frequency (F0) of 150 Hz while the upper note’s F0 ranged from 180 to 188 Hz (2 Hz spacing between adjacent tokens). Thus, the musical interval continuum spanned a minor (token 1) to major third (token 5). The minor-major third continuum was selected because these intervals occur frequently in Western tonal music and connote typical valence of “sadness” and “happiness”, respectively, and are therefore easily described to participants unfamiliar with music-theoretic labels (Bidelman & Walker, 2017). Moreover, without training, nonmusicians perceive musical intervals in a continuous mode indicating they are initially heard non-categorically (Bidelman & Walker, 2017, 2019; Burns & Ward, 1978; Howard et al., 1992; Locke & Kellar, 1973; Siegel & Siegel, 1977; Zatorre & Halpern, 1979).

### Procedure

Participants were seated comfortably in an electroacoustically shielded booth. Stimuli were presented binaurally through ER-2 insert earphones (Etymotic Research) at ∼81±1 dB SPL. Stimulus presentation was controlled by MATLAB routed through a TDT RP2 interface (Tucker Davis Technologies). Categorization was assessed in a pre- and post-test phase.

Following brief task orientation (∼2-3 exemplars), tokens of the continuum were randomly presented on each trial. Participants were instructed to label the sound they heard as either “minor” or “major” via keyboard button press as fast and accurately as possible. The interstimulus interval was 400-600 ms (jittered in 20 ms steps) following the listener’s response. No feedback was provided during the pre- or post-test. To reduce fatigue, participants were offered a break before and after the training phase.

#### Training paradigm

Participants in the learning group underwent a 20-min identification training between the pre- and post-test phases (all performed in a single ∼3 hr period). Training consisted of 500 trials, 250 presentations each of the minor and major 3^rd^ exemplars (i.e., tokens 1 and 5), spread evenly across 10 blocks^1^. Feedback was provided to improve accuracy and efficiency of auditory category learning (Yi & Chandrasekaran, 2016). The training procedure was conducted using E-Prime 2.0 (PST, Inc.).

### EEG acquisition and preprocessing

EEG data were recorded using a Synamps RT amplifier (Compumedics Neuroscan) from 64 sintered Ag/AgCl electrodes at 10-10 scalp locations and referenced online to a sensor placed ∼1 cm posterior to Cz. Impedances were <10 kΩ. Recordings were digitized at a sampling rate of 500 Hz. Preprocessing was completed in BESA Research (v7.1; BESA GmbH). Continuous data were re-referenced offline to the common average reference, epoched from - 200-800 ms, filtered from 1-30 Hz (4^th^-order Butterworth filter), baselined to the prestimulus interval, and averaged across trials to compute ERPs for each token per electrode.

### MRI segmentation and volumetrics

12 out of 19 learning group participants returned on a separate day for structural MRI scanning. 3D T1-weighted anatomical volumes were acquired on a Siemens 1.5T Symphony TIM scanner (tfl3d1 GR/IR sequence; TR = 2000 ms, TE = 3.26 ms, inversion time = 900 ms, phase encoding steps = 341, flip angle = 8°, FOV = 256 x 256 acquisition matrix, 1.0 mm axial slices). Scanning was conducted at the Semmes Murphey Neurology Clinic (Memphis, TN). All MRI T1-weighted images were initially registered to MNI ICBM 152 T1 weighted atlas with 1 x 1 x 1 mm^3^ isometric voxel size using affine transformation. The inverse transformation matrix was computed and applied to the brain mask in atlas space to create brain mask specific for each subject for skull removal (Evans et al., 1993). An LPBA40 T1 weighted atlas with 2 x 2 x 2 mm^3^ voxel size was then used to register the images and remove the cerebellum using the atlas cerebrum mask and following the same process as above (Shattuck et al., 2008). After skull removal and cerebrum extraction, an AAL3 T1 weighted atlas with 1 x 1 x 1 mm^3^ voxel size that provides parcellation of a large number of brain regions was used for extracting gray matter volume in certain regions of interest (ROIs) for each participant (Rolls et al., 2020).

### Data analysis

#### Behavioral data

Identification curves were fit with a two-parameter sigmoid function *P* = 1/[1 + *e*^-β1(*x*-β0)^], where *P* describes the proportion of trials identified as major, *x* is the step number along the stimulus continuum, β_0_ is the locus of transition along the sigmoid (i.e., categorical boundary), and β_1_ is the slope of the logistic fit. Larger β_1_ values reflect steeper psychometric functions and therefore better musical interval categorization performance. β_1_ slopes were square root transformed improve normality and homogeneity of variance. Reaction times (RTs) were computed as the listeners’ median response latency for the ambiguous (i.e., token 3) and prototypical tokens (i.e., mean[tokens 1 & 5]; see *ERP data*), after excluding outliers outside 250-2500 ms (Bidelman, Moreno, et al., 2013; Bidelman & Walker, 2017; Mankel, Barber, et al., 2020). As an index of training success, accuracy was calculated in the learning group as the average percent correct identification across all training trials.

#### ERP data

We analyzed a subset of electrodes from a frontocentral cluster (mean of F1, Fz, F2, FC1, FCz, FC2) where categorical effects in the auditory ERPs are most prominent at the scalp (Bidelman & Lee, 2015; Bidelman, Moreno, et al., 2013; Bidelman & Walker, 2017; Bidelman et al., 2014). Peak latencies and amplitudes were quantified for P1(40-80 ms), N1 (70-130 ms), and P2 (140-200 ms). The mean amplitude was also measured for slow wave activity between 300-500 ms, given prior work suggesting rapid auditory learning effects in this later time frame (Alain et al., 2010; Alain et al., 2007).

We also quantified neural responses at T7 and T8 to assess hemispheric lateralization. For these analyses, we computed difference waves derived between the ambiguous and prototypical tokens (ΔERP = mean[tokens 1 & 5] – token 3) for both the pre- and post-test (Bidelman, 2015; Bidelman & Walker, 2017; Mankel, Barber, et al., 2020). Larger ΔERP values indicate stronger differentiation of category ambiguous from category prototype sounds and thus reflect the degree of “neural categorization” in each hemisphere.

#### MRI data

Each participant’s MRI images were registered to the AAL3 atlas, ROI masks were transformed to subject space, and ROI volumes were then calculated (cm^3^). Atlas registration was confirmed using SPM12 toolbox in MATLAB (Penny et al., 2011). Cortical thickness was examined using a diffeomorphic registration based cortical thickness (DiReCT) measure (Das et al., 2009). We used the OASIS atlas (Marcus et al., 2009) for the computation of cortical thickness because it provides four brain segmentation priors for parcellating cerebrospinal fluid (CSF), cortical gray matter, white matter, and deep gray matter. 3D cortical thickness maps for each subject were computed based on these priors. Thickness maps were then multiplied with the AAL3 atlas (converted to subject space) to compute the cortical thickness of each brain region mapped to their corresponding labels. Finally, the mean, standard deviation, and range of the cortical thickness measurements along with the surface area and volume of the cortical regions were computed for each ROI. Volumetrics were normalized to each participant’s total intracranial brain volume to control for artificial differences across individuals (e.g., head size; Whitwell et al., 2001). To test for hemispheric differences specific to auditory neuroanatomic measures, we restricted ROI analysis to bilateral Heschl’s gyrus (PAC; Brodmann 41). MRI post-processing was performed using in-house scripts coded in Python (http://www.python.org).

### Statistical analysis

ERPs were analyzed using GLME mixed-effects regression models in SAS (Proc GLIMMIX; v9.4, SAS Institute, Inc.) with subjects as a random factor and fixed effects of training phase (two levels: pretest vs. posttest), token (two levels: tokens 1&5 vs. 3) and behavioral performance (identification slopes or training accuracy; continuous measures). We also included the interaction of phase and behavioral performance to investigate whether brain-behavior correspondences change after training. Similar models were used to analyze the behavioral and MRI data. Analyses on the individual groups alone included main and interaction effects of identification slopes or training accuracy (learning group only), training phase, and stimulus token. We used a backward selection procedure to remove nonsignificant variables and report final model results throughout. Post hoc multiple comparisons were corrected using Tukey adjustments. Identification function slopes were square-root transformed to reduce right skewness. Demographic variables were analyzed using Wilcoxon-Mann-Whitney and Fischer’s exact tests due to non-normality. An *a priori* significance level was set at α = 0.05. Conditional studentized residuals, Cook’s D, and covariance ratios were used to identify and exclude influential outliers.

## RESULTS

### Training results

Behavioral training outcomes are plotted in **Figure 1**. On average, participants in the learning group improved 10-15% in accuracy (**Fig. 1A**; *F*_9,158_ = 2.05, *p* = 0.038) and exhibited faster RTs (**Fig. 1B**; *F*_9,149_ = 3.22, *p* = 0.001) over the course of training. Training was highly effective; most individuals averaged >80-90% identification accuracy across the 10 blocks (i.e., the approximate performance of a musician on the same task; data not shown). N=5 “nonlearners” had training accuracies that remained near chance performance; these individuals were removed for subsequent analysis. Post hoc analyses revealed RTs became faster following the second training block (all *p*’s < 0.05; block 1 vs. 2 *p* = 0.052). Similarly, listeners’ identification was more accurate starting at the 9^th^ training block compared to the first block (block 9 vs. 1: *t*_158_ = 3.44, *p* = 0.025; block 10 vs.1: *t*_158_ = 3.40, *p* = 0.028).

**Figure 1:**
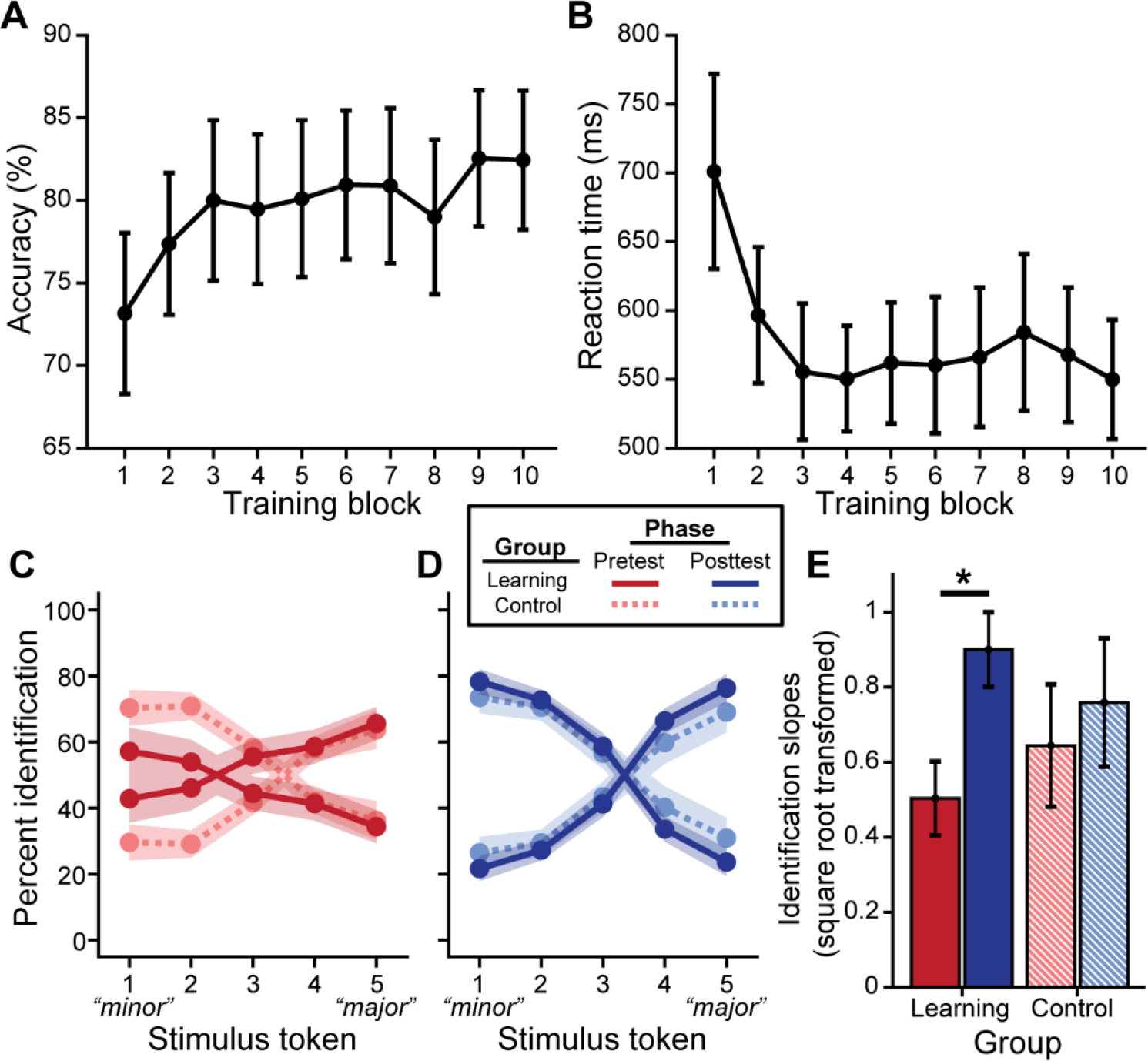
Behavioral categorization improves following rapid auditory training. Brief major/minor categorization training yields an increase in accuracy (A) and decrease in reaction time (B) across blocks. Pretest (C) and posttest (D) psychometric identification functions show stronger categorization for musical intervals after training for the learning group (excluding data from n=5 nonlearners); performance was identical pre- to post-test for control listeners (E). Error bars/shading = +/- 1 SE. \**p* < 0.05.

### Behavioral categorization following training

We then assessed training training-related improvements in categorization via listeners’ identification of the musical interval continuum. We found a group x session interaction for identification slopes (*F*_1,26_ = 4.93, *p* = 0.035). Importantly, control and learning groups did not differ at pretest (**Fig. 1C**; *t*_26_ = -0.14, *p* = 0.48), suggesting common baseline categorization. Critically, post hoc analyses revealed that identification slopes were steeper at posttest for successful learners (**Fig. 1D-E**; *t*_26_ = 4.42, *p* < 0.001), whereas performance remain static in the control group (*t*_26_ = 4.42, *p* = 0.21). For learners, in addition to training gains (main effect of phase: *F*_1,13_ = 11.65, *p* = 0.005), achieving better accuracy during training was associated with steeper identification functions overall (*F*_1,13_ = 8.58, *p* = 0.012). Similarly, RTs showed a group x phase interaction (*F*_1,78_ = 3.98, *p* = 0.050). Whereas the control group achieved faster RTs at posttest (*t*_78_ = -3.64, *p* < 0.001), RTs remained constant in the learning group (*t*_78_ = -0.73, *p* = 0.47).

### Electrophysiological results

ERP waveforms are shown per group and experimental phase in **Figure 2** (pooling all tokens). For the learning group, we found a training accuracy x phase interaction in P2 (*F*_1,39_ = 5.77, *p* = 0.021) and P1 amplitudes (*F*_1,39_ = 11.29, *p* = 0.002); better performance during training was associated with decreased amplitudes in the posttest but not the pretest (P2 posttest: *t*_39_ = - 3.41, *p* = 0.010; P1 posttest: *t*_39_ = -2.32, *p* = 0.010). All other ERP comparisons with training accuracy were not significant.

**Figure 2:**
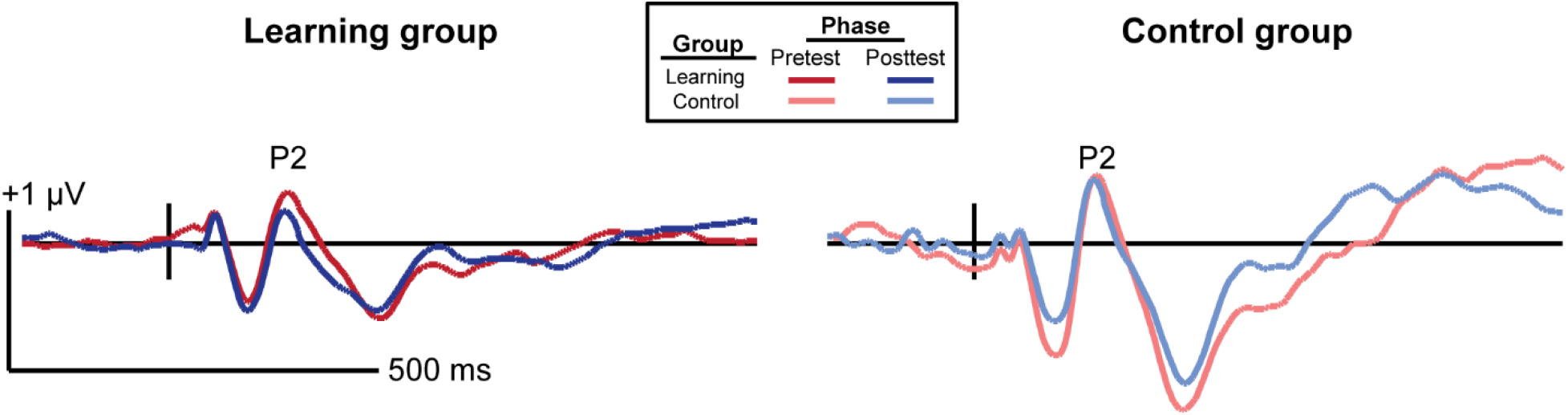
Grand average ERP waveforms collapsed across all tokens from the frontocentral electrode cluster (mean F1, Fz, F2, FC1, FCz, FC2). The learning group (left) underwent brief identification training whereas the control group (right) did not. The tick mark represents t=0 (stimulus onset).

In learners, we found an identification slopes x phase interaction for P2 amplitudes (*F*_1,38_= 4.16, *p* = 0.048); steeper (i.e., more categorical) posttest identification slopes were associated with a decrease in neural activity after training (**Fig. 3A**). Main effects of slope (*F*_1,39_ = 8.46, *p* = 0.006) and phase (*F*_1,39_ = 6.26, *p*=0.017) were also found for the slow wave (300-500 ms). Critically, these brain-behavior relationships were specific to learners and were not observed in the control group (**Fig. 3B**; all *p’s* > 0.05).

**Figure 3:**
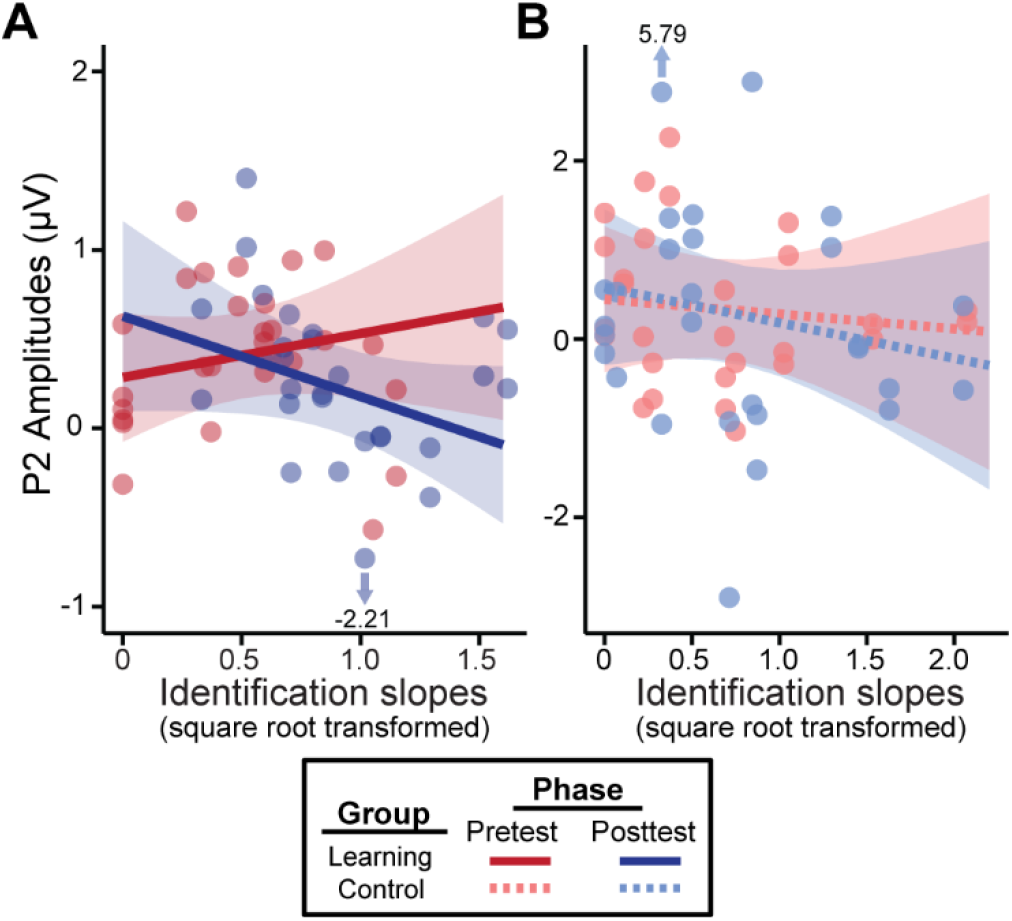
Neural amplitudes scale with behavioral outcomes in the learning group (A) but not the control group (B). Better posttest categorization (i.e., steeper identification slopes) is associated with a decrease in P2 amplitudes. Data points indicate individual subjects (collapsed across tokens 1 & 5 and 3). Arrows/values mark outliers (which did not alter results). Shading = 95% CI.

Hemispheric asymmetries were assessed via difference waveforms (i.e., mean[tokens 1 & 5] vs. 3) indexing the degree of categorization contained in neural responses. This analysis focused on electrodes T7 and T8 located over the left and right temporal lobes, respectively. We used a running paired t-test to evaluate training effects in a point-by-point manner across the ERP time courses (BESA Statistics, v2; **Fig. 4**). This revealed that in learners, category differentiation was modulated by learning 112-356 ms after stimulus onset. Training effects were most prominent over electrode T8 (right hemisphere; **Fig. 4B**). Guided by these results, we then extracted average amplitudes within this time window and ran a three-way mixed model ANOVA (group, identification slopes, training phase). The group x slope interaction was significant for electrode T8 (*F*_1,23_ = 7.86, *p* = 0.010). Post hoc analyses revealed that for learners, steeper identification slopes predicted larger (i.e., more categorical) responses over the right hemisphere (*t*_23_ = 0.59, *p* = 0.021). This brain-behavior relationship was not observed in controls nor over the left hemisphere (*p’s* > 0.05). These data reveal a right hemisphere bias in neural mechanisms supporting category learning of musical sounds.

**Figure 4:**
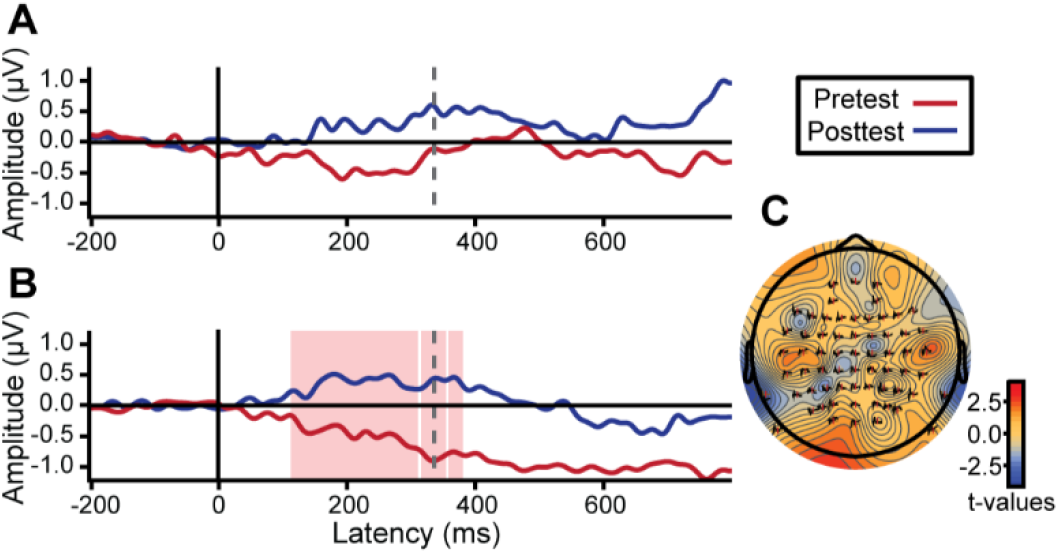
Neuroplastic changes following auditory categorical learning of music intervals is biased toward right hemisphere. Only data for the learning group is shown. (A-B) Difference waves (i.e., mean[token 1/5] – token 3) indexing categorical neural coding. An increase in neural categorization after training occurs over right (B; electrode T8) but not left hemisphere (A; electrode T7). Shaded region indicates a significant session effect (*p* < 0.05). (C) Topographic statistical map at t = 336 ms (dotted gray line in A & B) where pre- to post-test changes in categorical coding is maximal.

### Neuroanatomical results

Having established that musical interval learning leads to functional lateralization, we next assessed whether preexisting structural asymmetries (i.e., gray matter volume, cortical thickness) of primary auditory cortex were also associated with successful category learning. Volumetric analyses revealed that gray matter volumes were larger on average in the right compared to left PAC (*t*_11_ = 12.36, *p* < 0.001). The interaction of phase and structural measures were not significant for identification slopes. However, phase was kept in the models to isolate the relationship between structural PAC measures and behavior after factoring out training effects (see section *3*.*2*). Smaller gray matter volumes in right PAC were associated with stronger categorization overall (*F*_1,11_ = 5.80, *p* = 0.035, after accounting for effects of phase) (**Fig. 5**). Meanwhile, thinner cortical thickness of left PAC corresponded to better identification slopes (*F*_1,11_ = 15.07, *p* = 0.003, after accounting for effects of phase). Cortical thicknesses and gray matter volumes did not correlate with each other for either right or left PAC suggesting these volumetrics provided independent measures of the anatomy (all *p’s* > 0.05). Taken together, these results indicate that preexisting differences in bilateral PAC structure predict individual categorization performance.

**Figure 5:**
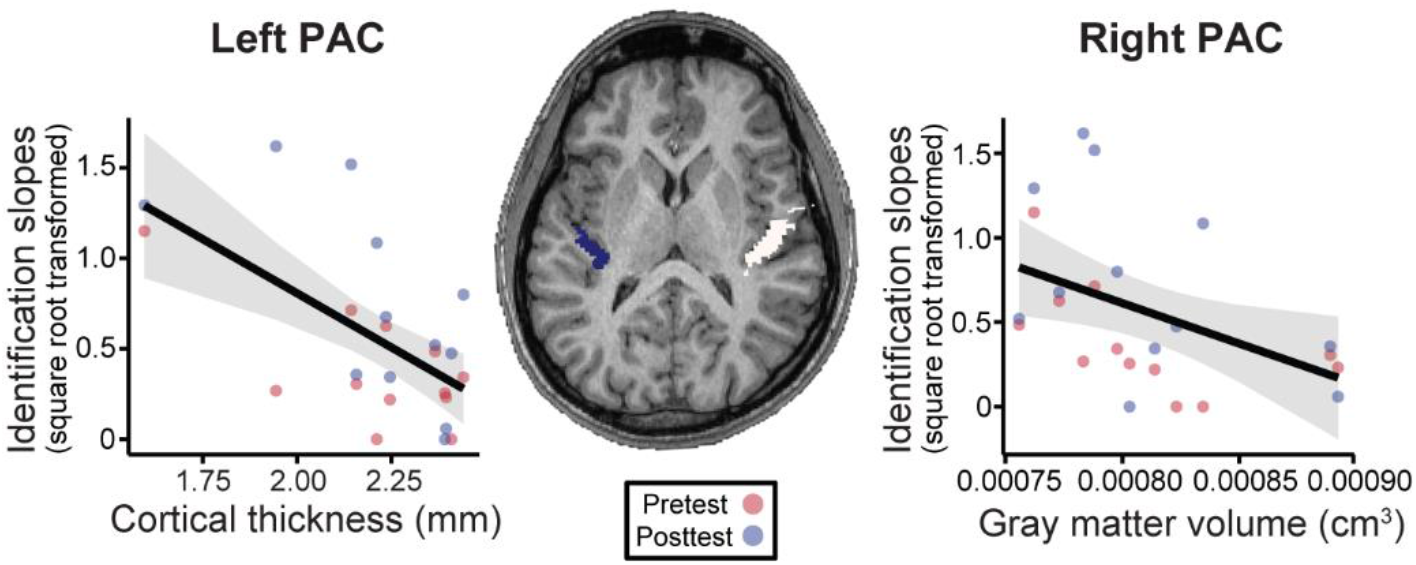
Neuroanatomical measures in primary auditory cortex (PAC) predict behavioral categorization performance in the learning group. (Left) In left HG, larger cortical thickness is associated with poorer categorization. (Right) Similarly, larger gray matter volumes (cm3) in right HG were associated with poorer behavioral categorization. (Center) MRI image from a representative subject with left and right HG shown in blue and white, respectively. Data points indicate individual subject identification slopes. Shading = 95% CI.

## DISCUSSION

By measuring multichannel EEGs and brain volumetrics during short term auditory category learning tasks, our data reveal three primary findings: (i) rapid label learning of non-speech sounds emerges very early in the brain (∼150-200 ms, P2 wave), (ii) ERP responses decrease with more successful learning suggesting more efficient neural processing (i.e., reduced amplitudes) after training; (iii) neuroplastic changes in categorizing musical sounds are stronger in right hemisphere where smaller and thinner auditory cortical regions predicted better categorization performance. Successful category learning is therefore characterized by increased functional efficiency of sensory processing layered on preexisting structural advantages within auditory cortex.

### Functional correlates of auditory category learning

Our data suggest category acquisition for non-speech sounds is associated with changes in ERP P2. The functional significance of P2 is still poorly understood (Crowley & Colrain, 2004). Experience-dependent neuroplasticity in P2 has been interpreted as reflecting enhanced perceptual encoding and/or auditory object representations (Bidelman & Lee, 2015; Bidelman et al., 2014; Garcia-Larrea et al., 1992; Ross et al., 2013; Shahin et al., 2003), improvements in the task acquisition process (Tremblay et al., 2014), reallocation of attentional resources (Alain et al., 2007), increased inhibition of task-irrelevant signals (Seppanen et al., 2012; Sheehan et al., 2005), or mere stimulus exposure (Ross et al., 2013; Sheehan et al., 2005). Here, we demonstrate early ERP waves including P1 (∼40-80 ms) as well as P2 (∼150-200 ms) closely scale with behavioral learning. Moreover, these neuroplastic effects are surprisingly fast, occurring rapidly within only 20 minutes of training. Our findings parallel visual category learning where changes in the visual-evoked N1 and late positive component signal successful learning (Perez-Gay Juarez et al., 2019). Our results also align with previous studies using various auditory training tasks including speech (Alain et al., 2010; Alain et al., 2007; Ben-David et al., 2011; Tremblay et al., 2001; Tremblay & Kraus, 2002; Tremblay et al., 2009) and nonspeech sounds (Atienza et al., 2002; Bosnyak et al., 2004; Tong et al., 2009; Wisniewski et al., 2020) suggesting P2 indexes auditory experience that reflects learning success and is not simply a product of the task acquisition process (cf. Tremblay et al., 2014) or repeated stimulus exposure (Ross et al., 2013; Ross & Tremblay, 2009; Sheehan et al., 2005). The lack of clear neural effects in control listeners further rules out exposure or repetition effect accounts of our data.

In this study, successful learning (i.e., both training accuracy and identification function slopes) was characterized by a *reduction* in ERP amplitudes after training. The specific direction of P2 modulations varies across experiments with some reporting an increase in evoked responses with learning (Atienza et al., 2002; Bosnyak et al., 2004; Carcagno & Plack, 2011; Ross et al., 2013; Sheehan et al., 2005; Tong et al., 2009; Tremblay et al., 2001; Wisniewski et al., 2020) and others a decrease (Alain et al., 2010; Ben-David et al., 2011; Zhang et al., 2005). As suggested by Alain et al. (2010), such discrepancies could be related to the task (e.g., active task vs. passive recording), the stimuli (e.g., speech vs. non-speech), the rate of learning among the participants, or even the rigor of training paradigm. Studies reporting enhanced P2 often included multiple days of training or recorded ERPs during passive listening (Atienza et al., 2002; Bosnyak et al., 2004; Ross et al., 2013; Seppanen et al., 2013; Tremblay et al., 2001; Wisniewski et al., 2020). Long-term auditory experiences (e.g., music training, tone language expertise) have also been associated with enhanced P2 during active sound categorization (Bidelman & Alain, 2015; Bidelman & Lee, 2015; Bidelman et al., 2014) as well as learning (Seppanen et al., 2012, 2013; Shahin et al., 2003). The ERP decreases we find in successful learners are highly consistent with single-session, rapid learning experiments demonstrating greater efficiency of sensory-evoked neural responses during active task engagement (Alain et al., 2010; Ben-David et al., 2011; Guenther et al., 2004; Perez-Gay Juarez et al., 2019; Sohoglu & Davis, 2016). Consequently, our results reinforce notions that the P2 is a biomarker of learning to classify auditory stimuli and map sounds to labels.

On the contrary, decreased neural activity might reflect other aspects of the task, including arousal and/or fatigue (Crowley & Colrain, 2004; Näätänen & Picton, 1987). However, decreased neural activity from these factors would have been expected in both groups due to the similar task constraints on all participants. If better learners simply sustain arousal more effectively through posttest, we would have also expected faster RTs. Rather, our data suggest decreases in activation meaningfully reflect music category learning, paralleling findings with speech (Guenther et al., 2004). Alternatively, given modulations in both P2 and slow wave activity, a separate but overlapping processing negativity within this timeframe cannot be ruled out. Negative processing components have been associated with early auditory selection and attention (Crowley & Colrain, 2004; Hillyard & Kutas, 1983; Näätänen & Picton, 1987) and may therefore be another target for learning processes.

### Hemispheric lateralization and music categorization

Our findings show that acquiring novel categories for musical intervals predominantly recruits neural resources from the right auditory cortex, complementing the left hemisphere bias reported for speech categorization (Alho et al., 2016; Chang et al., 2010; Desai et al., 2008; Golestani & Zatorre, 2004; Liebenthal et al., 2005; Liebenthal et al., 2010; Myers et al., 2009; Zatorre et al., 1992). Specifically, we observed enhanced neural categorization over the right hemisphere in more successful learners. Gray matter volumetrics in right PAC also predicted behavioral categorization abilities. These findings support long-standing notions about lateralization for speech vs. music categorization in the brain (Alho et al., 2016; Bidelman & Walker, 2019; Bouton et al., 2018; Chang et al., 2010; Desai et al., 2008; Klein & Zatorre, 2011, 2015; Liebenthal et al., 2010; Mankel, Barber, et al., 2020; Zatorre et al., 1992). Superior music categorization in both trained musicians (Bidelman & Walker, 2019; Klein & Zatorre, 2011, 2015) as well as musically adept non-musicians (Mankel, Barber, et al., 2020) has been associated with right temporal lobe functions. We thus provide new evidence that even brief, 20-minute identification training is sufficient to induce neural reorganization in the right hemisphere circuity that subserves auditory sensory coding and classification of musical stimuli.

### Neuroanatomical correlates of auditory category learning

Our MRI results demonstrate that individual variation in structural measures (gray matter volume, cortical thickness) also predict behavioral categorization performance beyond mere training effects. Brain structure is influenced by genetic, epigenetic, and experiential factors (Zatorre et al., 2012). Thus, it is often difficult to know whether anatomical differences are innate or experience-driven, but structural measures are presumed to be more stable and less plastic than functional responses (e.g., ERPs) (Golestani, 2012). Anatomical variability in auditory cortex has been related to learning rate and attainment for foreign speech sounds (Golestani et al., 2007), linguistic pitch patterns (Wong et al., 2008), and melody discrimination (Foster & Zatorre, 2010) as well as native speech categorization (Fuhrmeister & Myers, 2021). Consistent with this prior work on speech, our findings suggest that individual differences in music category perception and functional plasticity are influenced by anatomical predispositions within auditory cortex—that is, a layering of both nature and nurture.

It is often assumed larger morphology within a particular brain area yields better computational efficiency (Kanai & Rees, 2011) (i.e., “bigger is better”). For example, faster, more successful learners of nonnative speech sounds show more voluminous primary auditory cortex and adjacent white matter in left hemisphere (Golestani et al., 2007; Golestani et al., 2002; Wong et al., 2008). Relatedly, expert listeners (i.e., musicians) show increased gray matter volumes and cortical thickness in PAC (Bermudez et al., 2009; de Manzano & Ullen, 2018; Gaser & Schlaug, 2003; Schneider et al., 2002; Seither-Preisler et al., 2014; Wengenroth et al., 2014). Instead, our data show the opposite pattern with regard to non-speech category learning. To our knowledge, only one study has shown correspondence between decreased gyrification in temporal regions and improved consistency in speech categorization behaviors (Fuhrmeister & Myers, 2021). Similarly, smaller gray matter volume in STG has been linked to improvements in speech and cognitive training (Maruyama et al., 2018; Takeuchi, Taki, Hashizume, et al., 2011; Takeuchi, Taki, Sassa, et al., 2011). Thus, it seems “less is more” with respect to the expanse of auditory anatomy and certain aspects of listening performance. However, future research is needed to clarify the relationships between macroscopic gray and white matter volumes measured by MRI, neuronal microstructures, and their behavioral correlates.

## Conclusion

We demonstrate that rapid auditory category learning of musical interval sounds is characterized by increased efficiency in sensory processing in bilateral, though predominantly right, auditory cortex. The relationship between better behavioral gains in identification performance and the ERPs corroborate P2 as an index of auditory experience and a biomarker for successful perceptual learning. The right hemisphere dominance supporting music category learning complements left hemisphere networks reported for speech categorization. These short-term functional changes can be superimposed on preexisting structural differences in bilateral auditory areas to impact individual categorization performance.

## Acknowledgements

Requests for data and materials should be directed to G.M.B. This work was supported by the National Institute on Deafness and Other Communication Disorders of the National Institutes of Health under award numbers F31DC019041 (K.M.) and R01DC016267 (G.M.B.).

## Footnotes

1 Two pilot subjects received 6 and 15 blocks of training, respectively, before settling on the final 10 block training regimen.

## Notes

### Competing Interest Statement

The authors have declared no competing interest.

